# Chromomycin A_5_ induces bonafide immunogenic cell death in metastatic melanoma

**DOI:** 10.1101/2021.09.18.460876

**Authors:** Katharine G. D. Florêncio, Evelline A. Edson, Francisco C. L. Pinto, Otília D. L. Pessoa, João Agostinho Machado-Neto, Diego V. Wilke

## Abstract

Some first-line cytotoxic chemotherapics, *e.g.* doxorubicin, paclitaxel and oxaliplatin, induce activation of the immune system through immunogenic cell death (ICD). Tumor cells undergoing ICD function as a vaccine, releasing damage-associated molecular patterns (DAMPs), which act as adjuvants, and neoantigens of the tumor are recognized as antigens. ICD induction is rare, however it yields better and long-lasting antitumor responses to chemotherapy. Advanced metastatic melanoma (AMM) is incurable for more than half of patients. The discovery of ICD inducers against AMM is an interesting drug discovery strategy with high translational potential. Here we evaluated ICD induction of four highly cytotoxic chromomycins A (CA_5-8_). B16-F10, a metastatic melanoma cell line, treated with CA_5-8_ and doxorubicin exhibited ICD features such as autophagy and apoptosis, externalization of calreticulin, and releasing of HMGB1. However, CA_5_-treated cells had the best profile, also inducing ATP release, ERp57 externalization, phosphorylation of eIF2α and altering expression of transcription of genes related to autophagy, endoplasmic reticulum stress, and apoptosis. Bonafide ICD induction by CA_5_ was confirmed by a C57BL/6 mice vaccination assay with CA_5_-treated cells. These findings support a high potential of CA_5_ as an anticancer candidate against AMM.

## 1. Introduction

Advanced metastatic melanoma (AMM) is the most aggressive skin cancer, and is a serious concern due to increasing incidence in recent decades. Chemotherapy with dacarbazine and temozolomide was the standard of care in metastatic melanoma until 2011, however with no benefit for overall survival (QUEIROLO et al., 2019). Immunological therapy with IL-2 induces long-lasting responses in a small subset of patients, albeit with a high rate of severe toxicities (ATKINS et al., 2016; QUEIROLO et al., 2019). Advances in AMM treatment have yielded better agents based on target therapies, such as BRAF and MEK inhibitors (KAKADIA et al., 2018) and immunological checkpoint inhibitors, e.g. anti-PD-L1, anti-PD-1 and anti-CTLA-4 antibodies (LARKIN et al., 2015; QUEIROLO et al., 2019). Despite these advances, more than half of patients still do not experience a satisfactory clinical response. Thus translational research should pay particular attention to the non-responders subset of patients, aiming to shift immune-cold tumors into immune-hot ones (QUEIROLO et al., 2019).

The immunogenic effect of first-line chemotherapy and some radiotherapy treatments have been revealed as the hidden ally which improves responses of patients (KEPP et al., 2014; KROEMER et al., 2013; VANMEERBEEK et al., 2020). Treatment with inducers of immunogenic cell death (ICD) produces a specific antitumor immunity, which potentiates therapeutic efficacy (GALLUZZI et al., 2017; GARG et al., 2014; KEPP et al., 2014). ICD is a rare type of regulated cell death characterized by the activation of the adaptive immune system in the presence of cell death antigens, especially from cancer cells. Only 5% of chemotherapeutics the arsenal approved by the Food and Drug Administration of USA for cancer treatment are validated ICD inducers. However, they are first-line agents in the clinic and among the most used in the world including anthracyclines, taxanes and oxaliplatin (KEPP; SENOVILLA; KROEMER, 2014). This cell demise occurs under strong cellular stress, including autophagy and endoplasmic reticulum (ER) stress, and release of a constellation of damage-associated molecular patterns (DAMPs), the signals required to recruit and activate immune cells, such as antigen-presenting cells (APCs) and lymphocytes. The most important DAMPs in ICD include ATP, high mobility box group-1 (HMBG1), a nuclear non-histone nuclear factor, chaperones, specially calreticulin, and heat-shock proteins 70 and 90, annexin A1, CXCL10 and type I interferon (KROEMER et al., 2013; RADOGNA; DICATO; DIEDERICH, 2019). In addition to *in vitro* evaluation of cell demise associated with cell stress and release of DAMPs, a vaccination assay is required to validate induction of ICD through an anti-tumor response *in vivo* (GALLUZZI et al., 2020; VANMEERBEEK et al., 2020). Clinical evidence also supports ICD as a sensitizer to PD-1/PD-L1 blockade (KEPP et al., 2019). Thus identification of ICD inducers against AMM is a promising strategy for early identification of anticancer candidates with high expectation of translational success.

Chromomycins and mithramycins are promising antitumor antibiotic aureolic acids (KORMANEC et al., 2020). In 1970, mithramycin was approved for use in testicular cancer (BROWN AND TORKELSON, 1995), and chronic and acute myeloid leukemia (SNYDER et al., 1991; DUTCHER et al., 1997). Recent studies have shown mithramycin inhibition of drug-resistant cancer-initiating stem cells, important players in the disease relapse (KORMANEC et al., 2020). Mithramycins also have been reported as an inhibitor of P-glycoprotein, a transmembrane efflux pump related to resistance to multiple drugs in cancer cells (TAGASHIRA et al, 2000).

Chromomycin A_3_ has antitumor activity, reversibly binding to minor DNA grooves by interacting with cytosine and guanine (CG)-rich DNA regions in the presence of Mg2+, preventing replication and transcription (SUKANYA CHAKRABARTI1; BHATTACHARYYA, 2000; CHAKRABORTY et al., 2014). GuimarÃes et al., (2014) showed chromomycin A_2_ induces autophagy in metastatic melanoma cells, MALME-3M. The pre-apoptotic autophagy is related to the immunogenic outcome of cancer cell death (MARTINS et al., 2012). Recently we obtained four cytotoxic dextrorotatory chromomycins A (CA_5_, CA_6_, CA_7_, and CA_8_) from the actinobacteria *Streptomyces* sp. BRA-384 which displayed IC_50_ values against five tumor cells from the pM to nM range (PINTO et al., 2019). CA_5_ binds to the transcription factor T-box 2 (TBX2), which also may be related to its antiproliferative effect and antimetastatic potential (SAHM et al., 2020). Because CA_5-8_ are highly cytotoxic (PINTO et al., 2019) and likely to induce autophagy of melanoma cells (GUIMARÃES et al., 2014), we hypothesized these compounds were bonafide ICD inducers. Here we investigated the induction of ICD by CA_5-8_ on an AMM model, which could mark the renaissance of chromomycins as promising anticancer agents.

## 2. Materials & Methods

### 2.1 Reagents

Chromomycins (CA_5_, CA_6_, CA_7_, and CA_8_) were obtained as previously described by Pinto et al. (2019). Doxorubicin and dimethyl sulfoxide (DMSO) were purchased from Sigma-Aldrich (Missouri, USA). All cytotoxic compounds were diluted in DMSO.

### 2.2 Cell culture

The murine metastatic melanoma B16-F10 cell line was purchased from Banco de células do Rio de Janeiro (Rio de Janeiro, Brazil) and cultured following the manufacturer’s instructions.

### 2.3 Animals

We used a total of 21 C57BL/6 mice (female, 18–20 g) 6-8 weeks-old, free of ecto and endoparasites, obtained from the animal house of the Federal University of Ceara, Brazil. Animals were housed in cages under a 12:12 h light-dark cycle (lights on at 6:00 a.m.) and food and water *ad libidum*. All animal handling procedures were performed following the Brazilian legislation for the use and care of laboratory animals (No 11.724/20080) after approval by the Animal Ethics Committee of the Federal University of Ceara (No 3000310818).

### 2.4 Antiproliferative assay

Sulforhodamine B (SRB) assay was performed as described by Skehan et al., 1990. CA_5-8_ 0.32 to 1000nM (CA5, CA6, CA7 and CA8 respectively), doxorubicin at 0.6μM (Dox) as positive control and DMSO (0.05%) as negative control were added to cells during 4h, 8h, 12h, 24h, 48h and 72h and antiproliferative effect evaluated after 72h. When the exposure time was < 72h, cells were washed and replaced by fresh media (see Fig. 1). Inhibition concentration mean (IC_50_), total growth inhibition (TGI), and lethal concentration mean (LC_50_) values were calculated from cell growth percentage normalized data through interpolation of nonlinear regression using GraphPad Prism v6 (GraphPad Software, Inc., San Diego, CA, USA).

**Figure 1.**
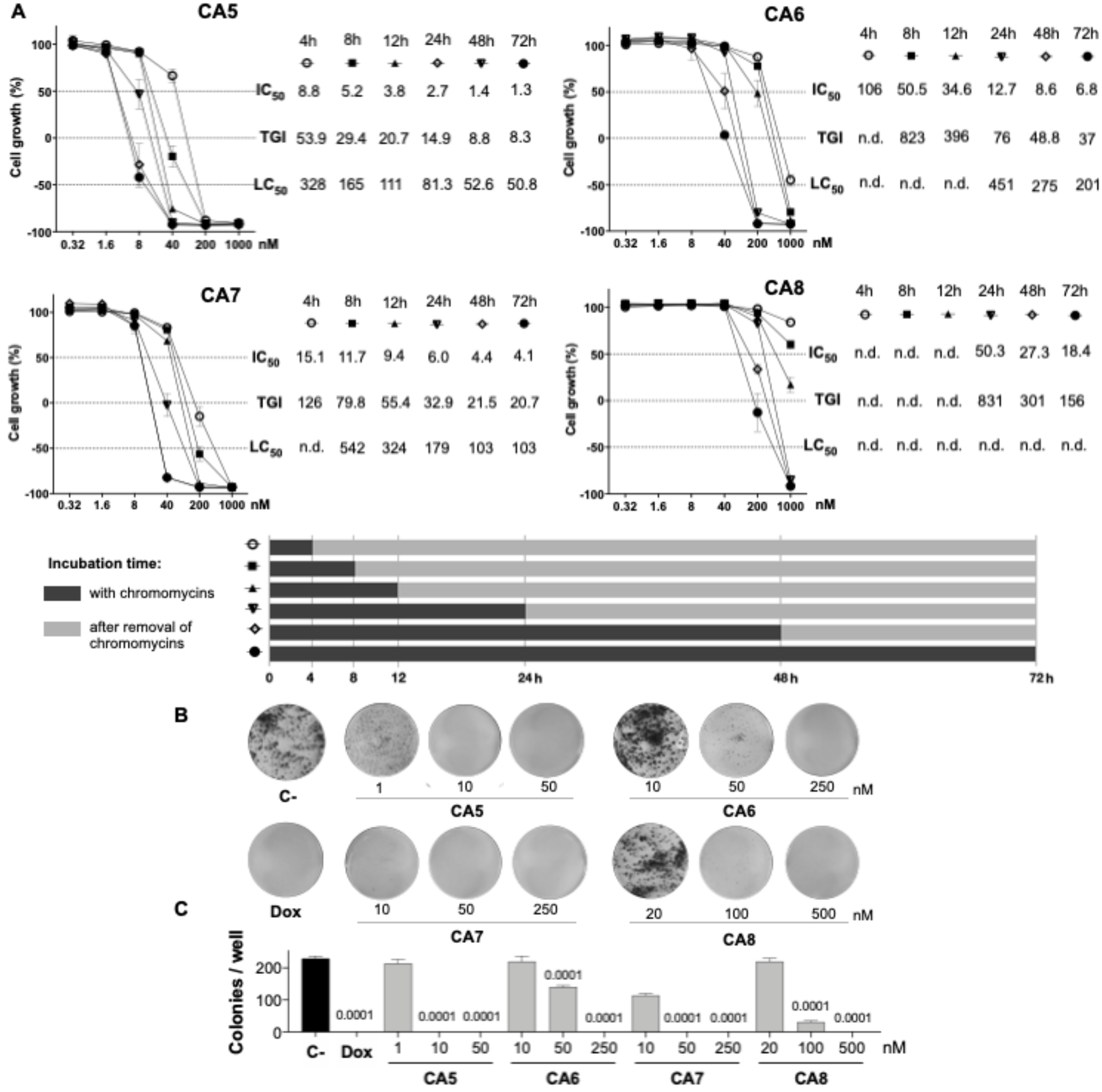
Antiproliferative profile of chromomycins A_5-8_ on metastatic melanoma B16-F10 cell line over time. **A**, Graphs showing the antiproliferative activity of CA_5-8_ after different exposure periods by the SRB assay. Inhibition concentration mean (IC_50_), total growth inhibition (TGI), and lethal concentration mean (LC_50_) values were obtained from interpolation of non-linear regression of normalized absorbance of 3 experiments performed in triplicate. **B**, Representative photos of the clonogenic assay, and **C**, Number of colonies represented as the mean ± standard deviation. 0.05% DMSO (C-) and 0.6 μM doxorubicin (Dox) were considered as a negative control and positive control, respectively. Statistical differences of treated groups versus C- are expressed as p values indicated above the columns of the groups.

### 2.5 Clonogenic assay

The colony-forming assay was performed according to Franken et al., 2006. Briefly, cells were seeded in 6-well plates at a density of 500 cells/well and exposed to CA_5-8_ (CA5, CA6, CA7, and CA8), doxorubicin (Dox) or 0.05% DMSO (C-) for 24 hours. After this period, the medium with cytotoxic agents or DMSO was replaced by a fresh medium, and plates were analyzed daily until the DMSO control reached a high density of individualized colonies (approximately 7 days). Cells were then washed with PBS and stained with violet crystal dye (0.5% crystal violet in methanol 50% and distilled water). Total individual colonies/well were counted under a stereoscopic microscope.

### 2.6 Cell treatment for immunogenic cell death investigation

Usually, 2.5 × 10^4^ cells/mL were seeded in 24-well plates for flow cytometry assays, 96-well for ATP assay, or in Petri dish plates (90 × 15 mm) for western blot and qPCR assays, and incubated for 24 h before the treatment. Then cells were exposed to CA_5_ at 0.1μM (CA5), CA_6_ at 0.25 μM (CA6), CA_7_ at 0.25 μM (CA7) and CA_8_ at 0.5 μM (CA8), doxorubicin at 0.6 μM (Dox), as the ICD inducer positive control, and 0.05% DMSO (C-), as the negative control, and incubated for 2 hours. After treatment, cells were collected and washed with PBS before all protocols described below.

### 2.7 Cell viability - flow cytometry

All flow cytometry assays were set to acquire 10,000 events excluding debris and doublets using a FACSVerse™ flow cytometer (BD Biosciences, San Diego, CA, USA) and FlowJo v10.6 software (Ashland, OR: Becton, Dickinson and Company) for data analysis. Cell suspensions were incubated with 2 μg/mL 4’,6-diamidine-2’-phenylindole dihydrochloride (DAPI, Sigma-Aldrich, Missouri, USA) for 10 minutes as the final step in all flow cytometry assays to distinguish membrane integrity and disruption, except for the acridine orange stain.

### 2.8 Acidic vesicular organelles (AVOs) staining with acridine orange (AO) – flow cytometry

Differential acridine orange (AO) staining is an assay that is strongly correlated with autolysosomes formation in the late step of autophagy. The AO assay was performed as described previously (THOMÉ et al., 2016). Briefly, cells were washed with PBS and stained with 1 μg/mL AO (Sigma-Aldrich, Missouri, USA) for 30 min in the dark at room temperature. Excitation of AO-stained cells with a 488 nm laser induces green fluorescence in whole cells and red fluorescence is produced in acidic vesicular organelles (AVOs) due to AO metachromasia. AVOs were gated in a region with an increased ratio of red fluorescence. The increase of cell granularity, although nonspecific, is a relevant cell stress alteration found in autophagy and ER stress. High cell granularity was gated in the high side scatter (SSC) region.

### 2.9 Calreticulin (CRT) externalization – flow cytometry

Cells were fixed with 0.25% paraformaldehyde in ice-cold PBS for 5 min and incubated with anti-CRT (Calreticulin (D3E6) XP Rabbit mAb, Cell Signaling Technology, Danvers, MA, USA, #12238) 1:300 for 40 minutes, in the dark at 4 °C. Cells were washed with a FACS buffer (FACS solution supplemented with 4% fetal calf serum) and incubated with anti-rabbit secondary antibody Alexa Fluor 488™ (#4412, Cell Signaling Technology, Danvers, MA, USA)(1:800) for 40 minutes, in the dark at 4 °C. Cells were centrifuged and resuspended in the FACS buffer for acquisition in the flow cytometer. The percentage of Ecto-CRT was counted as Low FSC / CRT^+^ population in the DAPI^−^ cells.

### 2.10 ERp57 (Ser51) externalization measurement – flow cytometry

Cells were fixed with 0.25% paraformaldehyde in ice-cold PBS for 5 min and incubated with anti-ERp57 (#A484, Rabbit mAb, Cell Signaling Technology, Danvers, MA, USA) 1:100 for 40 minutes, in the dark at 4 °C. Cells were washed with a FACS buffer (FACS solution supplemented with 4% fetal calf serum) and incubated with anti-rabbit secondary antibody conjugated with Alexa Fluor 488™ (1:400) for 40 minutes, in the dark at 4 °C. Cells were centrifuged and resuspended in the FACS buffer for acquisition in the flow cytometer. The mean fluorescence intensity (MFI) related to Ecto-ERp57 was evaluated in the DAPI^−^ population.

### 2.11 Evaluation of nuclear HMGB1 - flow cytometry

Cell membranes were permeabilized with 0.1% TritonX 100 solution for 5 minutes. Then cells were washed with PBS and incubated with anti-HMGB1 conjugated with phycoerythrin (PE) (# 651403, PE anti-HMGB1; Biolegend, San Diego, CA, USA) for 40 min, in the dark at 4 °C. After incubation, cells were washed, resuspended in the FACS buffer, and acquired in the flow cytometer. The HMGB1 release was estimated indirectly by the MFI decreasing in cells exposed to doxorubicin and CA_5-8_ (GOMEZ-CADENA et al., 2016).

### 2.12 Evaluation of eIF2α and P-eIF2α – flow cytometry

Cell membranes were permeabilized with 0.1% TritonX 100 solution for 5 minutes. Then cells were washed with PBS and incubated with anti-eI2Fα (#9722, Cell Signaling Technology, Danvers, MA, USA) or anti-P-eIF2α (phospho S51) (#9721, Cell Signaling Technology, Danvers, MA, USA). Antibodies (1:100) were incubated for 40 min at 4 °C. Subsequently, these cells were washed and incubated with the secondary antibody conjugated with Alexa Fluor 488 (Biolegend, San Diego, CA, USA) (1:400) for 40 minutes, in the dark at 4 °C. After cell wash and resuspension in the FACS buffer, the data were acquired by flow cytometry. MFI of eIF2α and P-eIF2α was measured in the DAPI^+^ region.

### 2.13 ATP release assay

ATP determination kit (#A22066, ThermoFisher Scientific, Inchinnan, UK) based on luciferin-luciferase conversion was carried out according to the manufacturer’s protocol. Briefly, the supernatant was centrifuged at 1200 rpm for 5 min and 10 μL of cleared supernatants of each condition were transferred to a 96-well plate for luminescence. Then, 90 μL of ATP mix reagent was added to each well. After incubation for 1 min at room temperature, we analyzed luminescence emission in the multimode microplate reader Cytation 3 (Biotek, Vermont, USA).

### 2.14 Western blot analysis

Total protein extraction was performed using a buffer containing 100 mM Tris (pH 7.6), 1% Triton X-100, 150 mM NaCl, 2 mM PMSF, 10 mM Na_3_VO_4_, 100 mM NaF, 10 mM Na_4_P_2_O_7_, and 4 mM EDTA. Equal amounts of protein were used from total extracts followed by SDS-PAGE, and Western blot analysis with the antibodies indicated, as previously described (LIPRERI DA SILVA et al., 2021) Antibodies against total and cleaved PARP1 (#9542), LC3BI/II (#2775), and α-tubulin (#2144) were obtained from Cell Signaling Technology (Danvers, MA, USA). Antibody against γ-H2AX (p-Histone H2A.X S139; sc-101696) was obtained from Santa Cruz Biotechnology (Santa Cruz, CA, USA). Antibody binding was revealed using a SuperSignalTM West Dura Extended Duration substrate system (Thermo Fisher Scientific) and a G: BOX Chemi XX6 gel document system (Syngene, Cambridge, UK).

### 2.15 Quantitative RT-PCR (qRT-PCR)

B16-F10 cells were seeded on cell culture dishes (90×15mm) and treated with 0.05% DMSO, chromomycin CA5 (0.1 μM), or doxorubicin (0.6 μM) for 24 h. Total RNA was obtained using TRIzol reagent (Thermo Fisher Scientific). cDNA was synthesized from 1 μg of RNA using a High-Capacity cDNA Reverse Transcription Kit (Thermo Fisher Scientific). Quantitative PCR (qPCR) was performed using a QuantStudio 3 Real-Time PCR System in conjunction with a SybrGreen System (Thermo Fisher Scientific) in conjunction with a SybrGreen System for the expression of *Atf4*, *Atf6*, *Atg5, Atg7, Bak1, Bad, Bax, Bcl2, Becn1, Carl, Hspa4, Hspa5 and Sqstm1* genes. *Actb* and *Hprt1* were used as reference genes. A negative ‘No Template Control’ was included for each primer pair. All procedures were performed according to the manufacturer’s instructions. Relative quantification values were calculated using the 2^-ΔΔCT^ equation (LIVAK; SCHMITTGEN, 2001). The heatmap was constructed using the multiple experiment viewer (MeV) 4.9.0 software (SAEED et al., 2003). The network analysis was performed using modulated genes by Dox or CA5 groups using the GeneMANIA tool (WARDE-FARLEY et al., 2010).

### 2.16 Vaccination assay

The vaccination assay was performed with a syngeneic mouse model as described by GOMEZ-CADENA et al., 2016 with modifications. A syngeneic mouse model (e.g., B16-F10 cell line in C57BL/6 mice), is an appropriate approach to study cancer therapy with a functional immune system. The vaccination was performed on day −7 when the mice of the 3 experimental groups (N=7 animals/group) received subcutaneously into the right axilla 200 μL of 0.9% saline solution (Saline), as a negative control, or 200μL of dying B16-F10 cells pre-exposed 24 h to 0.1 μM CA_5_ (CA5) and 0.6 μM doxorubicin (Dox), as a positive control. The dying cells were obtained after exposure to cytotoxic agents for 24 h, then harvested, washed with PBS twice, and resuspended in PBS with 1.8 × 10^5^ cells/200 μL. No additional adjuvants were added. Seven days later, on day 0, mice of Saline, CA5, and Dox groups were challenged with an injection of 1.0 × 10^5^ viable B16-F10 cells in 200 μL of PBS into the left armpit subcutaneously. The size of the tumors was measured at days 12, 15, and 17 with digital calipers. The tumor volume was calculated using the following formula: tumor volume (in mm^3^) = [(width)^2^ × length] / 2.

### 2.17 Statistical analysis

All statistics were performed in GraphPad Prism v6 (GraphPad Software, LLC, San Diego, CA, USA). Shapiro-Wilk was used to test the normal distribution of results. Data are expressed as means ± standard deviation of the mean (SD). Comparisons between C-and treated groups were performed using the One-Way Analysis of Variance (ANOVA) followed by Dunnet’s post-test for parametric data and using Kruskal-Wallis followed by Dunn’s post-test for nonparametric data. A *p*<0.05 value was considered significant.

## 3. Results

### 3.1 Chromomycins A_5-8_ are highly cytotoxic at multiple time exposures

Initially, we performed concentration-effect curves with CA_5-8_ varying time exposure to determine their antiproliferative profile against metastatic melanoma B16-F10 cells (Fig.1). CA_5_ and CA_7_ were the most potent compounds, depicting low to mid nM cytostatic and cytotoxic effects respectively even at short time exposures (Fig. 1A). In addition, CA_6_ also displayed a similar profile at longer exposures. CA_8_ induced a potent cytostatic effect, however, it failed to show cytotoxicity at the nM range. Additionally, CA_5-8_ inhibited colony formation at the nM range of B16-F10 cells incubated for 24 h (Fig. 1B and C).

### 3.2 Chromomycins A_5-8_ induce apoptosis

Immunogenic cell death (ICD) determination requires confirmation of early and late apoptotic features (GALLUZZI et al., 2020). B16-F10 cells depicted typical apoptosis features after exposure to CA_5-8_ such as cell shrinkage (Fig. 2A and Fig. 2B) and PARP1 cleavage (Fig. 2D). Notably, CA_5_ induced intense PARP1 cleavage compared to the other chromomycins, while the Dox group was similar to C-. All treated groups increased (p<0.0001) the cell death population subset (Fig. 2B and C). Similar to PARP1 cleavage, CA_5-8_ also induced an increase of **γ**-H2AX levels, and Dox did not elicit this marker of DNA damage (Fig. 2D).

**Figure 2.**
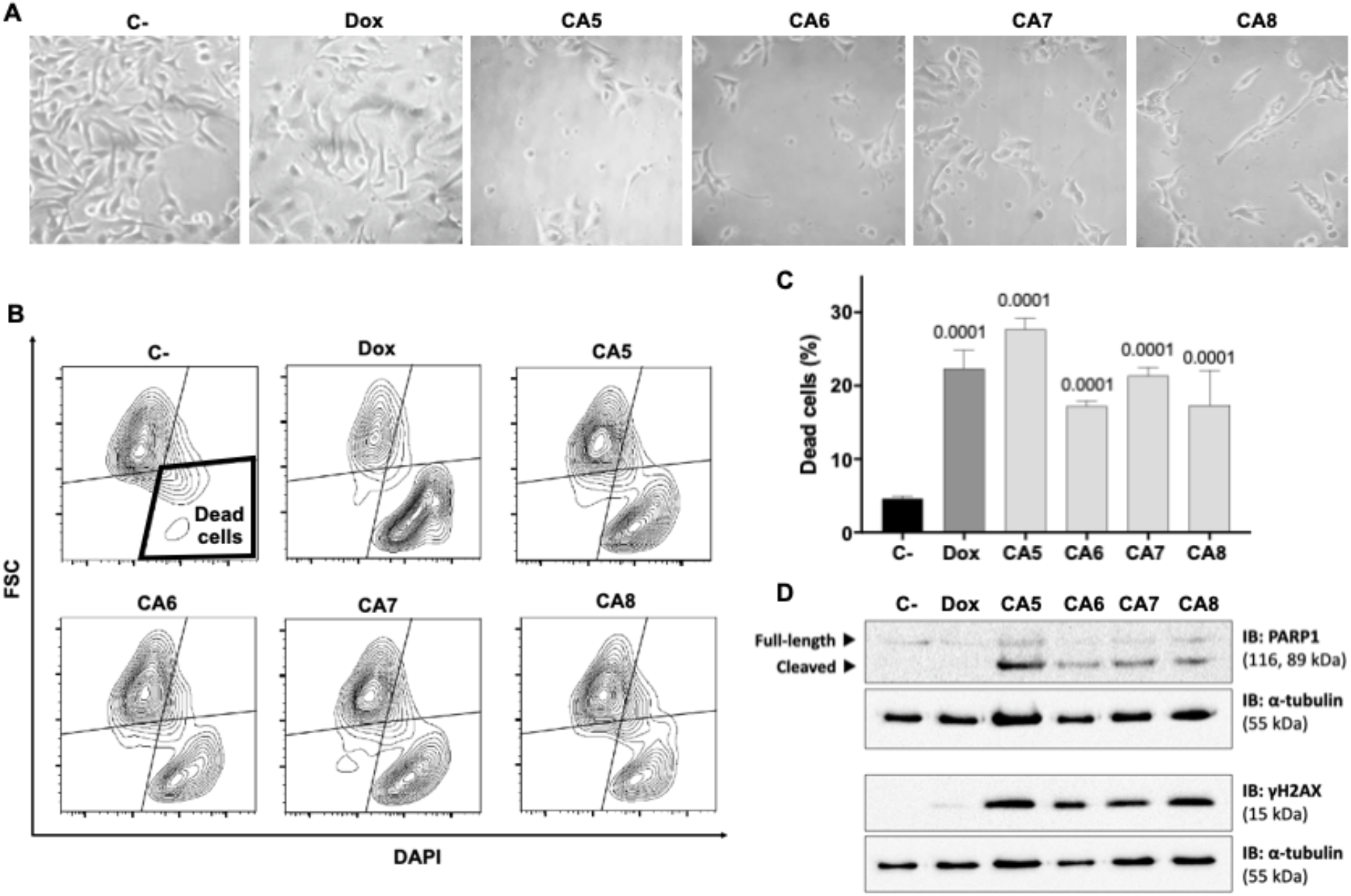
Chromomycins A_5-8_ induce cellular morphological changes and cell death. B16-F10 melanoma cells were treated with DMSO (C-), as a negative control, doxorubicin (Dox), as a positive control, and CA_5-8_ (CA5-8) for 24 h. **A**, Phase contrast photomicrographs (200x). **B**, Representative contour plot graphs of the cell death subpopulation gated in the low forward scatter (FSC) and DAPI+ region by flow cytometry and **C,** the respective graph depicting the percentage of dead cells. Data presented as mean ± standard deviation of 3 independent experiments performed in triplicate. The *p* values of C- compared to the treated groups are above each treated group. **D**, Expression of cleaved PARP1 and **γ**H2AX obtained by Western blot. Values associated with test proteins were normalized to standard α-tubulin for the relative expression measure.

### 3.3 Chromomycins induce autophagy

ICD inducers elicit cell stress associated with cell demise. Autophagy and endoplasmic reticulum (ER) stress are phenotypic changes observed quite often with ICD (GARG; AGOSTINIS, 2014). CA_5-8_ and Dox significantly increased the granularity of B16-F10 cells (Fig. 3A and B). Additionally, CA_5-8_ and Dox increased cells with acidic vesicular organelles (AVOs) (Fig. 3C and D), and LC3B I/II levels as well (Fig. 3E). These data confirm the autophagy induction by chromomycins on B16-F10 cells.

**Figure 3.**
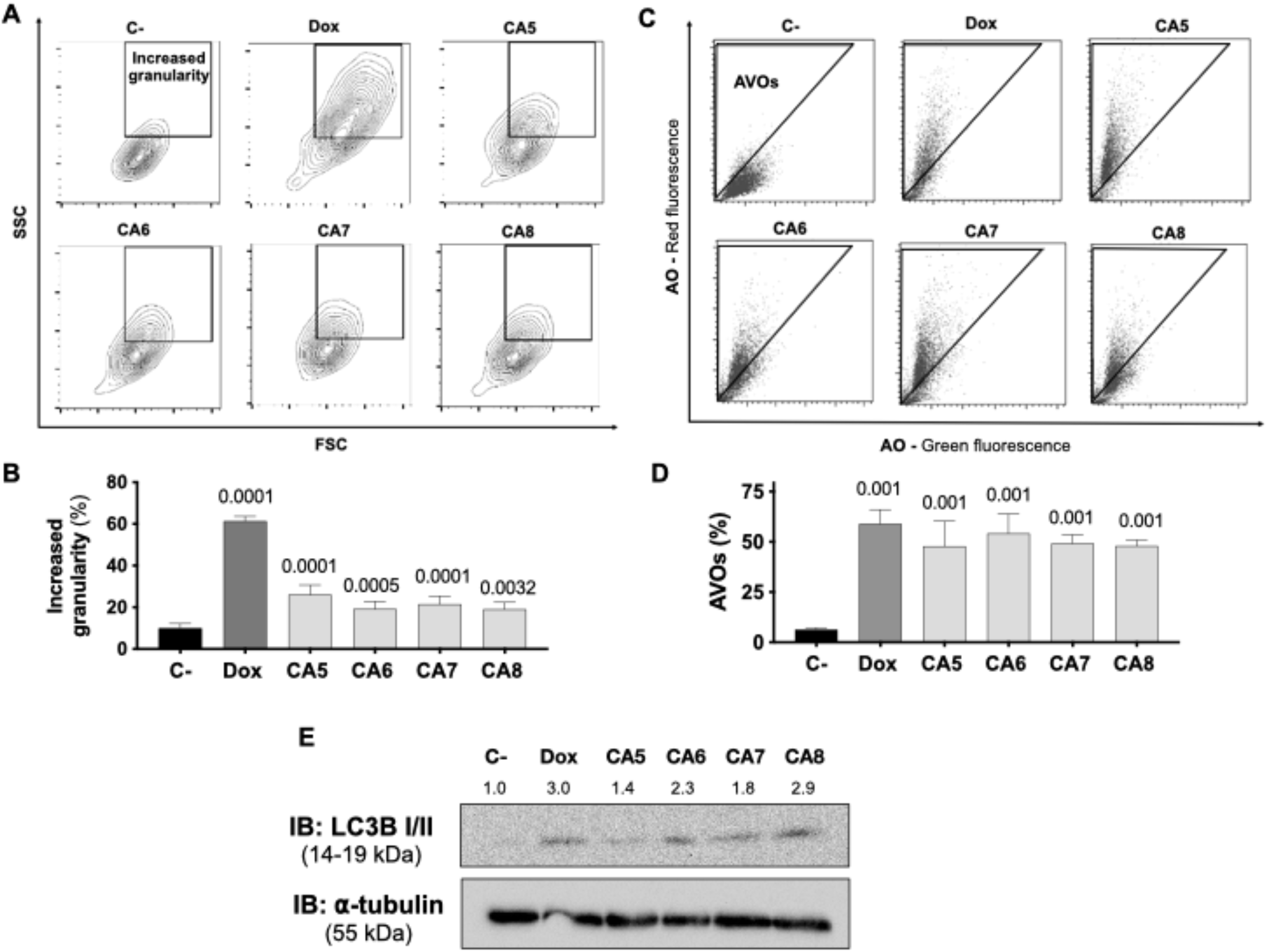
Chromomycins A_5-8_ induce autophagy. B16-F10 melanoma cells were treated with DMSO (C-), as a negative control, doxorubicin (Dox), as a positive control, and CA_5-8_ (CA5-8) for 24 h. **A**, Representative contour plot graphs of cells with granularity gated on high side scatter (SSC) and average forward scatter (FSC) region by flow cytometry and **B**, the graph depicting the percentage of high granularity cells. **C**, Representative dot plots of acridine orange (AO) staining. Acidic vesicular organelles (AVOs) were gated on the region with increased red fluorescence by flow cytometry and **D**, the graph showing the percentage of AVOs. Data presented in graphs as mean ± standard deviation of 3 independent experiments performed in triplicate. The *p* values of C- compared to the treated groups are above each treated group. **E**, Expression of LC3B I/II protein with values normalized by standard α-tubulin for the relative expression measure.

### 3.4 Chromomycins induce the release of ICD related DAMPs

The regulated cell death of B16-F10 exposed to CA_5-8_ associated with cell stress is a minimal requirement observed in ICD (Fig. 2 and Fig 3). However, the immunogenicity of ICD depends on the damage-associated molecular patterns (DAMPs) releasing. Several DAMPs related to ICD have been reported so far. Despite this, only a few examples are highly recurrent in ICD, such as secretion of ATP and HMBG1, a nuclear non-histone nuclear factor, and externalization to the plasma membrane of CRT, a lumenal chaperone of the ER (KROEMER et al., 2013; RADOGNA; DICATO; DIEDERICH, 2019; GALLUZZI et al., 2020). We observed nuclear HMGB1 decreasing in B16-F10 cells incubated with CA_5-8_ and Dox (Fig. 4A and B). ATP levels increased significantly, compared to C-, in supernatants of CA_5_ exposed cells, while Dox and CA_6-7_ did not change the level of this DAMP (Fig. 4C). Additionally, CA_5-8_ induced CRT externalization in the shrunken cells subpopulation (Fig. 4D, E, and F) and Dox as well. ATP and HMGB1 act as classic DAMPs, with chemoattractant and activation roles on APCs as ligands of purinergic receptors (P2Y2 and P2X7) and tool-like receptor 4 respectively. The ecto-CRT is a phagocytic signal recognized by CD91 that promotes antigen presentation of tumor neoantigens in presence of HMGB1 activation (GALLUZZI et al., 2020; GARG et al., 2012).

**Figure 4.**
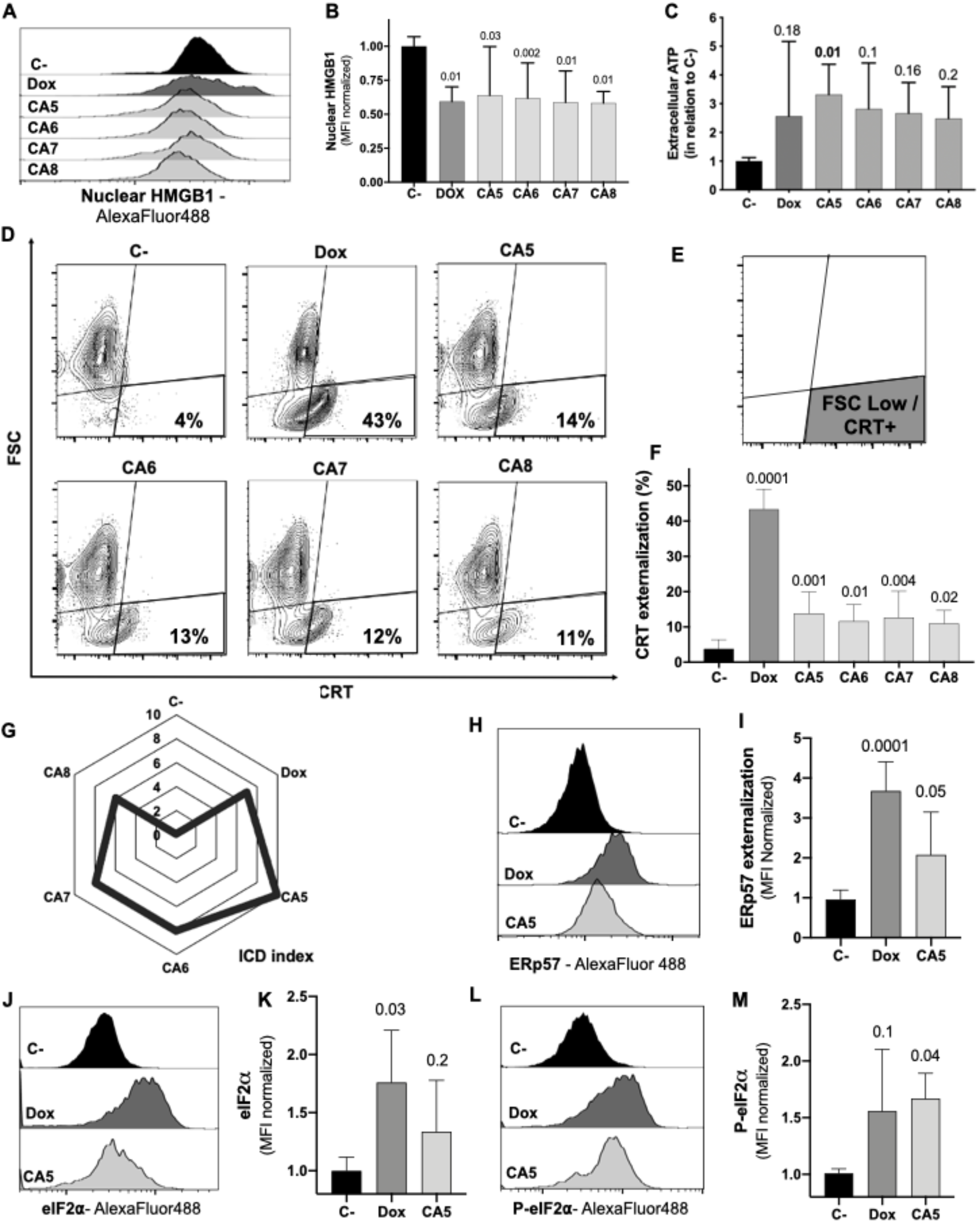
Chromomycins A_5-8_ induce the release of immunogenic cell death DAMPs. B16-F10 melanoma cells were treated with DMSO (C-), as a negative control, doxorubicin (Dox), as a positive control, and CA_5-8_ (CA5-8) for 24 h. **A**, Representative histograms of nuclear HMGB1 evaluated by flow cytometry and **B,** graph depicting normalized median fluorescence intensity (MFI) of nuclear HMGB1. **C**, Normalized extracellular ATP levels measured by luminescence. **D**, Representative contour plot graphs of calreticulin (CRT) vs cell size (forward scatter, FSC) by flow cytometry, **E**, Illustration of the gated region of CRT+ cells with low FSC and **F**, graph depicting the percentage of CRT+ cells. **G**, Radar graph of the index of immunogenic cell death (ICD index). Details of ICD index are described in SI.1. **H**, Representative histograms of ERp57 evaluated by flow cytometry and **I**, graph depicting normalized MFI of ERp57. **J**, Representative histograms of eIF2α evaluated by flow cytometry and **K**, graph depicting normalized MFI of eIF2a. **L**, Representative histograms of cells with eIF2α phosphorylated at serine 51 (P-eIF2α) evaluated by flow cytometry and **M**, graph depicting normalized MFI of P-eIF2a. Data presented in graphs as mean ± standard deviation of 3 independent experiments performed in triplicate. The *p* values of C- compared to the treated groups are above each treated group.

ICD is a complex phenomenon in which multiple phenotypic changes are required to allow the proper immune system activation. In order to compare the putative ICD potential among compounds used in our pipeline, we scored a total of seven parameters related to cell stress, cell death, and DAMPs from assays performed with CA_5-8_ and Dox as well (SI.1). The sum of scores for each compound was considered an ICD index, which aided us to understand the overall immunogenic potential profile of compounds tested in this study. CA_5_ depicted the highest ICD index followed by CA_6_ and CA_7_, Dox and CA_8_ respectively (Fig. 4G). From this point, we focused on further analyses of the effects of CA_5_.

The co-externalization of ERp57 with CRT is the actual “eat me” signal for phagocytosis by the APCs with an activation outcome. The B16-F10 cells exposed to CA_5_ and Dox presented ERp57 externalization (Fig. 4H and I). ER stress is an important ICD driver related to externalization of CRT and ERp57, and could initiate autophagy and apoptosis as well. The eIF2α is an ER stress protein involved in ICD. Dox-treated cells induced an increase of eI2Fα (Fig. 4J and K), while CA_5_ treatment did not change the level of this protein. Nevertheless, the phosphorylation of eI2Fα at Ser51 is the crucial ICD signaling (BEZU et al., 2018; HUMEAU et al., 2020). Cells incubated with CA_5_ elicited a significant activation of this protein (Fig. 4L and M). Curiously, Dox did not increase P-eIF2a levels despite the increase of the overall levels of eIF2a.

### 3.5 CA_5_ impacts gene expression related to autophagy, ER stress, and apoptosis

To obtain new insights into the molecular mechanisms involved in the response of B16-F10 cells to CA_5_, we investigated the expression of 13 genes related to autophagy, apoptosis, and ER stress by quantitative RT-PCR. A total of 7 of out 13 genes was significantly modulated by CA5 treatment (6 downregulated [*Atf4, Atf6, Hspa4, Hspa5, Atg5,* and *Sqstm1*] and 1 upregulated [*Benc1*], all p<0.05), while 5 genes were significantly modulated by Dox treatment (4 downregulated [*Atf4, Hspa4, Hspa5,* and *Atg5*] and 1 upregulated [*Bax*] all p<0.05) in B16-F10 cells. Of note, treatment with CA5, but nor Dox, significantly increased the *Becn1/Bcl2* and *Bad/Bcl2* ratios (Fig. 5A-B). The network analysis indicates that CA5 induces more complex relationships, which effectively interconnects the processes of apoptosis, autophagy, and ER stress (Fig. 5C).

**Figure 5.**
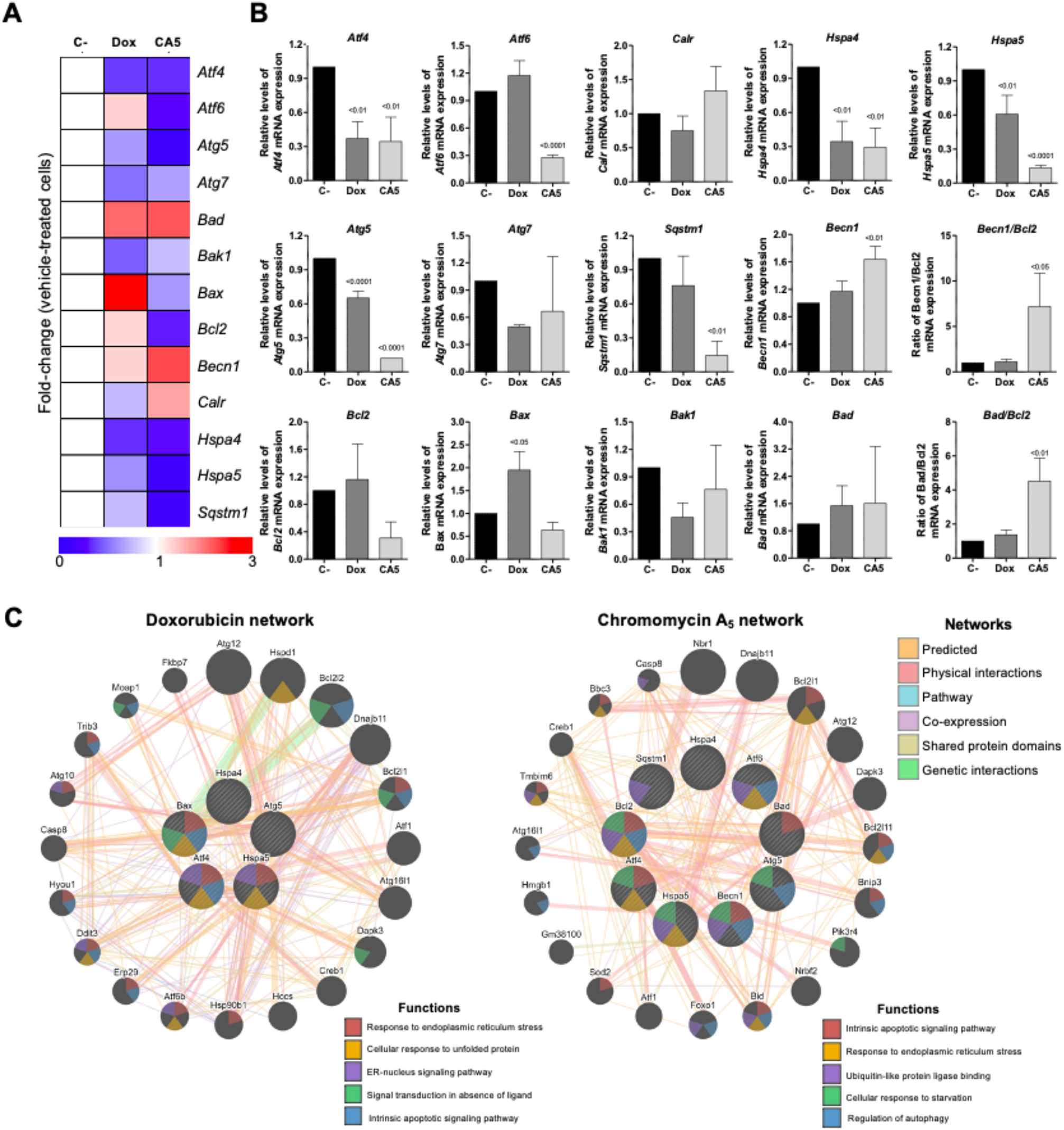
Chromomycin A_5_ modulates transcription genes related to endoplasmic reticulum (ER) stress, autophagy, and cell death. Quantitative RT-PCR was performed for 13 selected genes related to ER stress (*Atf4*, *Atf6*, *Calr*, *Hspa4*, and *Hspa5*); autophagy (*Atg5*, *Atg7*, *Becn1*, and *Sqstm1*); and apoptosis (*Bad*, *Bak1*, *Bax*, and *Bcl2*). **A**, Heatmap illustrating all selected genes in B16-F10 upon a vehicle, Dox (0.6 μM) or CA5 (0.1 μM) exposure for 24 h. The data are presented as fold-change of the vehicle-treated cells. Downregulated and upregulated genes are illustrated in blue and red, respectively. **B,** The comparison of selected genes are presented in bar graphs and the *p* values are indicated. **C,** Network analysis for genes modulated significantly by Dox or CA5 constructed using the GeneMANIA database (https://genemania.org/). The upregulated and downregulated genes in the quantitative RT-PCR are illustrated as strikethrough circles, and the interacting genes included by the software modeling are indicated by strikethrough strands. The main interactions between genes are indicated by colored lines and the five main cellular processes are described in the Figure.

### 3.6 Vaccination of mice with B16-F10 dying cells exposed to CA_5_

The vaccination assay using cells dying triggered by a cytotoxic agent is the gold standard technique for ICD confirmation (KEPP et al., 2014). CA_5_-exposed cells injected 7 days before the challenge with B16-F10 viable cells developed a significant tumor growth control protection (Fig. 6). This vaccination effect was not observed with mice of the Dox group. It is worth highlighting that the dying cells from CA_5_ and Dox groups were washed before injection in a saline solution.

**Figure 6.**
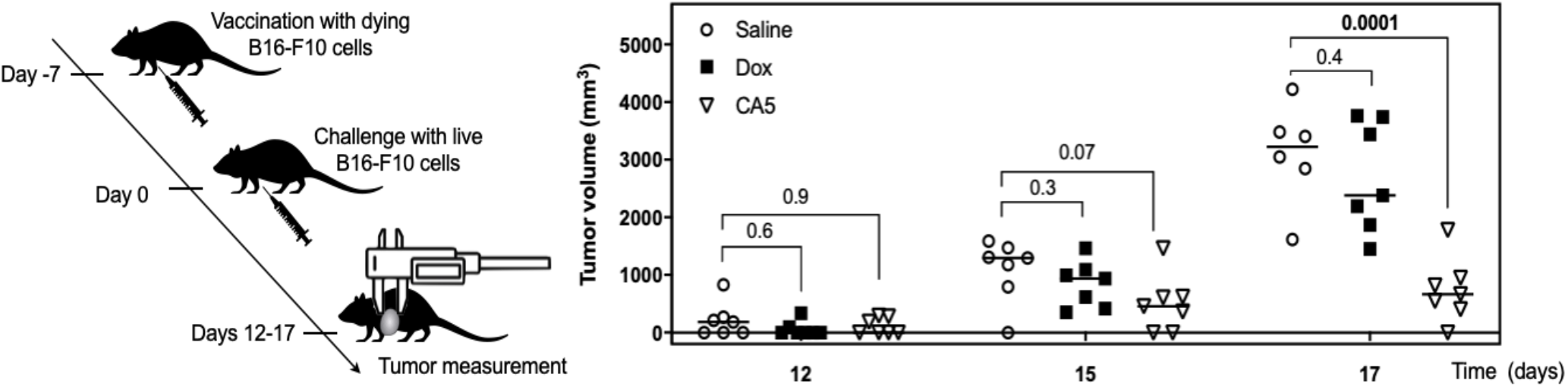
Vaccination with dying B16-F10 cells exposed to chromomycin A_5_ confers mice resistance against live B16-F10 cells. At day −7, 1.8 × 10^5^ cells pre-incubated for 24 h with 0.1 μM CA_5_ (CA5) and 0.6 μM doxorubicin (Dox) were injected subcutaneously in the right armpit of mice. 7 days after vaccination (day 0) mice were challenged with 1 × 10^5^ live B16-F10 cells in the left armpit. The mice of the negative control group (Saline) were injected with saline solution at −7 day and live B16-F10 cells at day 0. N= 7 mice/group. Tumor growth was monitored until day 17. Differences between groups are expressed as *p* values indicated above the compared groups.

## 4. Discussion

AMM lacks a therapeutic option to convert immune-cold into immune-hot tumors to improve the clinical response of the patients who do not respond to the current arsenal available including immunotherapy (BONAVENTURA et al., 2019). The identification of ICD inducers could fill this need by combining two useful effects at once, direct cytotoxicity against tumor cells and release of immunological activating signals. This type of regulated cell death allows the proper activation of the immune system, which in your turn, eliminates tumor cells resistant to chemotherapy. This mechanism is related to more effective and long-lasting responses (ZITVOGEL et al, 2008; ZITVOGEL et al., 2010; VANMEERBEEK et al., 2020). Herein we investigated the ICD induction of four chromomycins obtained from the marine bacterium *Streptomyces* sp. BRA-384 against metastatic melanoma B16-F10 cells.

Initially, the cytostatic and cytotoxic profiles of CA_5-8_ were investigated with increasing time exposure, and it was observed a time-dependent effect in the nM range. Notably, CA_5_ and CA_7_ depicted cytotoxicity in low time-exposure of 4 h and 8 h respectively (Fig. 1A). Additionally, CA_5-8_ completely inhibited colony formation of tumor cells after 24 h incubation (Fig. 1B). These data highlighted a favorable cytotoxic feature of chromomycins as anticancer compounds, which must achieve therapeutic plasma levels in a short time window due to toxicity.

The most reliable approach to the initial identification of ICD still consists in performing multiple phenotypic assays to evaluate autophagy, apoptosis, and releasing of DAMPs on treated cells *in vitro* (GALLUZZI et al, 2020). In general, cells treated with CA_5-8_ and Dox depicted some ICD features such as apoptosis (Fig. 2), autophagy (Fig. 3) and externalization of CRT, and releasing of HMGB1 (Fig. 4B, and F). Notably, CA_5_-treated cells depicted the most consistent ICD profile, filling all phenotypic features investigated so far, followed by CA_6_ and CA_7_, Dox and CA_8_ (Fig. 4G). Dox failed to increase cleaved PARP1 on B16-F10 cells, an apoptosis marker (KAUFMANN et al., 1993; TEWARI et al., 1995) and Dox and CA_6-8_ all failed to release ATP, an essential DAMP involved in the ICD (KEPP et al., 2014; HUMEAU et al., 2019; VULTAGGIO-POMA; SARTI; VIRGILIO, 2020). CA_5_ induced ATP release, ERp57 externalization, and phosphorylation of eIF2α (Fig. 4C, I and M respectively). Dox increased eIF2α levels, without significant phosphorylation at serine 51 (Fig. 4K and M). Analyses of anticancer ICD inducers revealed eIF2α phosphorylation mediated by eIF2α kinase-3 (EIF2AK3), but no other signs of ER stress are related to CRT exposure (PANARETAKIS et al., 2009; BEZU et al., 2018; HUMEAU et al., 2020). Furthermore, machine-learning approaches revealed eIF2α phosphorylation as the sole ER stress response relevant to the algorithm with downstream consequences including CRT exposure, stress granule formation, and autophagy induction (BEZU et al., 2018; HUMEAU et al., 2020). Although Dox did not induce a significant increase of phosphorylation of eIF2α, it induced CRT externalization (Fig. 4F) and increased the high granularity population (Fig. 3A and B) and autophagy (Fig. 3C-E). This conflicting data could be explained as a masking effect of the increase eIF2α in Dox-treated cells (Fig. 4K), which could produce biologically relevant phosphorylation of eIF2α as detected by increased ecto-CRT and autophagy.

CA_5_ and doxorubicin altered expression of transcription of 13 selected genes related to autophagy, ER stress, and apoptosis. However, CA_5_-treated cells changed most of the genes evaluated and increased the Becn1/Bcl2 and Bad/Bcl2 ratios (Fig. 5A-B) and depicted a wider interconnected network among apoptosis, autophagy, and ER stress than doxorubicin-treated cells (Fig. 5C). It is worth highlighting the cellular response to starvation along with ER stress and apoptosis, as a putative indication of a more intense stress response on CA_5_-treated cells in comparison to Dox. Activating transcription factors *Atf4* and *Atf6* were downregulated on CA5 cells (Fig. 5B). Dox treatment also decreased *Atf4*, however did not alter *Atf6*. ICD inducers, such as anthracyclines, enhance phosphorylation of eIF2α, but fail to stimulate other ER stress signs including the transcriptional activation of activating transcription factor 4 (ATF4) and the proteolytic cleavage of activating transcription factor 6 (ATF6) (BEZU et al., 2018; HUMEAU et al., 2020). The cellular changes induced by CA_5_ are summarized in the Fig. 7 along with the expected effect on dendritic cells.

**Figure 7.**
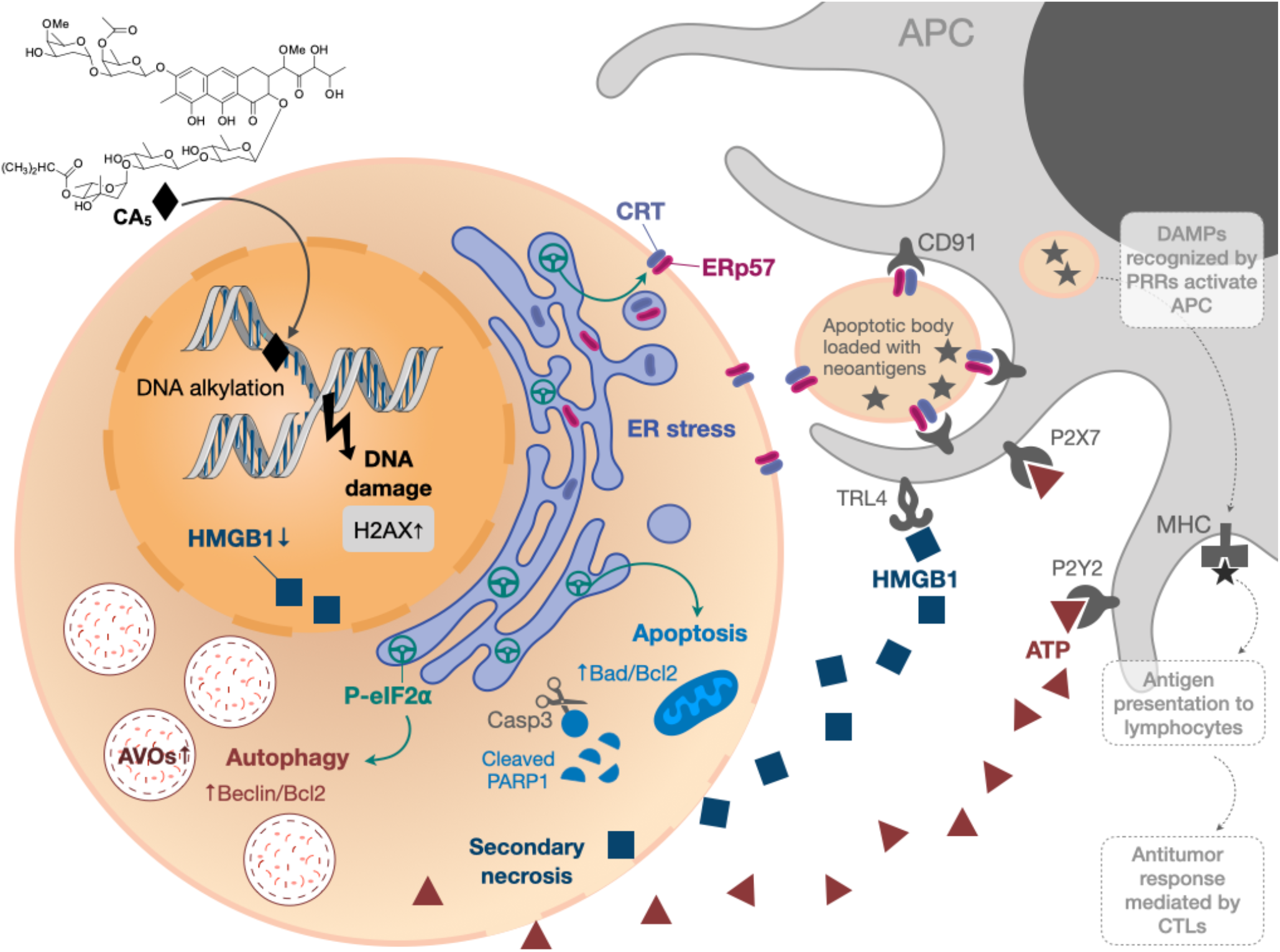
Overview of immunogenic cell death (ICD) induced by chromomycin A_5_ (CA_5_) in metastatic melanoma. B16-F10 cells treated with CA5 displayed cell stress and cell death along with release of damage-associated molecular patterns (DAMPs) which are supposed to activate antigen presenting cells (APC) and generating an effective cellular immune response. The phosphorylation of eIF2α due to endoplasmic reticulum (ER) stress drives crucial immunogenic events, as externalization of the “eat me” signals calreticulin (CRT) and ERp57, autophagy and apoptosis. AVOs, acidic vesicular organelles; Casp3, caspase 3; PRRs, pattern recognition receptors; CTLs, cytotoxic T lymphocytes.

The vaccination assay is the gold standard method to confirm ICD, due to its complex spatio-temporal nature. Dying cells undergoing bonafide ICD must effectively recruit and activate both APCs and lymphocytes without any external adjuvants. The vaccination efficacy is evaluated by challenging mice with live cells of the same lineage (KEPP et al., 2014; HUMEAU et al., 2019; VANMEERBEEK et al., 2020). C57BL/6 mice vaccinated with CA_5_-treated cells controlled tumor growth efficiently (Fig. 6). At day 17 the mean tumor volume of mice of the CA5 group was significantly lower (p=0.0001) than the mean tumor volume of the Saline group. Actually, animals from the CA5 group showed only 20% of the mean saline tumor volume, and one animal did not develop a tumor at all. This result confirms CA_5_ as a bonafide ICD inducer. Tumors of the Dox group did not show a significant difference from the negative control (p = 0.4). Gomez-Cadena et al. (2016) reported significant tumor control of C57L/6 mice vaccinated with Dox-treated B16-F10 cells. In their study, the cell treatment with doxorubicin was longer (48 h), and caspase 3 activation confirmed apoptosis induction. However, the ATP levels in supernatants of Dox-treated cells did not increase either, similar to our results as we found in the present study.

Fine-tuning *in vitro* conditions to confirm the induction of ICD is challenging and some studies could fail to demonstrate it depending on the histological origin of cells and time and concentrations of exposure used as well (SUKKURWALA et al., 2014). Most ICD inducers, such as doxorubicin, oxaliplatin, bortezomib and vinca alkaloids, were identified using tumor cell lines from different tumor origins of clinical practice (MENGER et al., 2012; SUKKURWALA et al., 2014). Although this approach generated robust knowledge about ICD inducers initially, it led to delayed identification of some important ICD inducer anticancer agents, including paclitaxel and cisplatin (LAU et al., 2020; SOLARI et al., 2020). Similarly, other chromomycins, including CA_7-8_ studied here, also induce ICD depending on experimental design; however further studies are needed to fully characterize ICD triggered by chromomycins. Additionally, the suboptimal results we obtained with melanoma cells exposed to doxorubicin, an important ICD inducer used in the treatment of several solid and hematological cancers (e.g. breast, ovary, prostate and multiple myeloma), also illustrates the challenge of identifying experimental conditions that trigger ICD.

A few chemotherapeutic agents are known to induce ICD, and they demonstrate remarkable clinical performance (KEPP; SENOVILLA; KROEMER, 2014), CA_5_ shows evidence of ICD and thus is a highly promising candidate for AMM and deserves further preclinical studies. It is also worth highlighting the supply as one important bottleneck to the preclinical and clinical development of pharmaceuticals (JIMENEZ et al., 2020). We obtained CA_5_ for this study using a sustainable and easily scalable technique (PINTO et al., 2019), and its supply for studies *in vivo* is quite feasible. In summary, we identified CA_5_ as a bonafide inducer of ICD in metastatic melanoma cells. Further *in vivo* studies with CA_5_ are necessary to evaluate antitumor activity, toxicity, and survival, as well as the effect on CA_5_ associated with immunotherapy.

## Abbreviations

AMM: advanced metastatic melanoma
AO: acridine orange
APC: antigen presenting cell
AVOs: acidic vesicular organelles
CRT: calreticulin
DAMPs: damage-associated molecular patterns
eIF2α: eukaryotic initiation factor 2α
ER: endoplasmic reticulum
HMGB1: high mobility group box-1
ICD: immunogenic cell death
PD-1: programmed death-1
PD-L1: programmed death-1 ligand

## Acknowledgements

We thank to Dr. Margo Haygood for reviewing the manuscript. This study was financed in part by the Coordenação de Aperfeiçoamento de Pessoal de Nível Superior - Brasil (CAPES) - Finance Code 001, Instituto Nacional de Ciência e Tecnologia (INCT BioNat-CNPq/FAPESP, No. 465637/2014-0) and Fundação de Amparo à Pequisa do Estado de São Paulo (2019/23864-7). The authors also thank the Multi-User Facility of Drug Research and Development Center of Federal University of Ceará for technical support.

## Author Contributions

**Katharine Gurgel Dias Florêncio**: Methodology, Validation, Writing - Original Draft, Formal analysis, Investigation, Visualization **Evelline Araújo Edson**: Methodology, Validation, Investigation, Formal analysis **Francisco das Chagas Lima Pinto**: Methodology, Validation, Investigation **Otília Deusdênia Loiola Pessoa**: Validation, review the manuscript **João Agostinho Machado Neto:** Methodology, Validation, Formal analysis, Investigation **Diego Veras Wilke**: Conceptualization, Funding acquisition, Supervision, Validation, Writing - Review & Editing.

## Supplementary information

**SI.1.** Immunogenic cell death (ICD) index. Seven parameters related to cell stress, cell death, and release of DAMPs from assays performed with CA_5-8_ and Dox were scored, based on statistical significance compared to negative control (C-) as 0 when p>0.05; 1 for cytotoxicity and autophagy parameters with p<0.05; and 2 for DAMPs releasing* with p<0.05. The sum of scores for each compound was considered the ICD index.

| Cellular phenotype |  | C- | Dox | CA5 | CA6 | CA7 | CA8 |
| --- | --- | --- | --- | --- | --- | --- | --- |
| Citotoxicity | Membrane disruption | 0 | 1 | 1 | 1 | 1 | 1 |
|  | Cleaved PARP1 | 0 | 0 | 1 | 1 | 1 | 1 |
| Autophagy | AVOs | 0 | 1 | 1 | 1 | 1 | 1 |
|  | LC3B I/II | 0 | 1 | 1 | 1 | 1 | 1 |
| DAMP releasing | **HMGB1** | 0 | 2 | 2 | 2 | 2 | 2 |
|  | **ATP** | 0 | 0 | 2 | 0 | 0 | 0 |
|  | **CRT** | 0 | 2 | 2 | 2 | 2 | 0 |
| **ICD index (sum)** |  | **0** | **7** | **10** | **8** | **8** | **6** |

\*The higher value of score for DAMPs releasing was arbitrarily chosen as an attempt to represent their critical importance on ICD induction.

